# Resolving *in vitro* heterogeneity of host functional responses to HCoV-229E via single-cell analyses

**DOI:** 10.64898/2026.04.17.719293

**Authors:** Anna H.C. Vlot, Vrinda Venu, Samantha Adikari, Eric M. Small, Cullen Roth, Kyle A. Sullivan, Shawn Starkenburg, Daniel A. Jacobson, Christina R. Steadman

## Abstract

Understanding host-pathogen interactions at the molecular level is essential to elucidating the mechanisms that govern immunity and pathogenesis. While diverse responses to viral infection have been documented, it remains unclear if this variability arises from the complex host milieu or represents an intrinsic property of the infection itself. To investigate this, we performed single-cell (sc) RNA sequencing analysis of infected MRC-5 fibroblast cells with human coronavirus 229E (HCoV-229E), which revealed pronounced variation in viral load across time points and biological replicates. While viral sequences were higher at 24 hours post-infection (hpi), more host genes were differentially expressed at 48 hpi, indicating dynamic responses during infection progression. Cells with higher viral load upregulated expression of unfolded protein response genes and NF-κB regulators, and downregulated expression of fibroblast identity and cell cycle genes, reflecting a shift toward a stress-adaptive, antiviral state. Network analyses identified NF-κB signaling as a central regulatory axis, with negative regulators overexpressed in high-viral-load clusters. Additionally, ATAC-seq analysis showed that regions of increased chromatin accessibility were enriched for NF-κB subunits, further supporting its role in transcriptional control. These findings demonstrate that even within ostensibly uniform cell cultures, viral infection induces distinct heterogeneous transcriptional and regulatory responses.

## INTRODUCTION

Host-pathogen interactions broadly encompass the diverse ways in which pathogens engage with their hosts, from entry and replication to survival and potential elimination. These interactions span scales from population transmission to cellular and molecular mechanisms, where pathogens adapt to evade host defenses and hosts counter with new immune strategies, often aided by therapeutic interventions. Several studies describe strong heterogeneity in immune responses, which influence infection outcomes^1^: thus, appropriate model systems are needed to disentangle intrinsic cellular variability from broad host impacts to understand infection persistence or clearance^2^. To this end, single-cell approaches have revealed differences in viral loads and corresponding immune responses across tissues and cell types^3,4^. Further, these studies report cell-type specific responses including differences among bystander and infected cells, changes across tissue type and infection progression, and differences among patients with variations in disease severity suggesting heterogeneity in host susceptibility^5–9^. While this heterogeneity in response to infection is often attributed to the complex host immune milieu, it may also arise from variability in viral progression and infection dynamics. Importantly, even in systems designed to reduce biological complexity, such as *in vitro* cell cultures, a uniform response to infection may be lacking. Indeed, there is growing appreciation for differential responses to viral infection among cells, but this has not been systematically tested. Thus, it remains unclear whether observed transcriptional heterogeneity arises solely from complex host tissue and immune environments or represents a fundamental property of viral infection itself.

Here, we consider that subtle cell-specific variability, even within homogeneous cell cultures, may yield insights about infectivity and host function. Understanding such variability can deepen our knowledge of disease propagation and help identify potential targets for novel therapeutics. To explore this potential variability, we employed the MRC-5 cell line, a fibroblast cell culture model system widely used to assess the efficacy of vaccines and in vaccine production to establish viral replication factories. We infected MRC-5 cells with human coronavirus 229E virus (HCoV-229E), an enveloped, positive-sense, single-stranded RNA virus which causes seasonal respiratory infections in humans. Notably, HCoV-229E infection induces heterogeneous outcomes ranging from mild illness in healthy adults to severe outcomes in infants, the elderly, and immunocompromised individuals^10–12^. Cell culture studies demonstrate that coronaviruses initiate reprogramming of host gene expression, including upregulation of immune response, stress adaptation, and protein folding, and downregulation of genes involved in cell cycle progression and differentiation^13^.

To assess transcriptional variability and potential heterogeneity of the MRC-5 cell culture system, we applied single-cell transcriptomics (scRNA-seq) to profile host functional responses across individual cells in response to HCoV-229E. Studies of related coronaviruses, including SARS-CoV-2, have assessed cell-to-cell variability in response to infection using scRNA-seq analyses, demonstrating cell-specific responses in patients^14–16^. Further, these reports suggest coordinated transcriptional and chromatin changes^17–19^; therefore, we integrated our scRNA-seq findings with bulk ATAC-seq analyses to unravel how chromatin remodeling contributes to the observed transcriptional reprogramming *in vitro*. Using predictive machine learning and *a priori* knowledge-driven, network-based analyses, we elucidate the regulatory mechanisms by which immune signaling, tissue remodeling, and cytokine responses drive heterogeneous transcriptional responses during HCoV-229E infection of MRC-5 cells. This integrated approach resolved cell-level variability in host responses and maps infection-mediated regulatory pathways to epigenomic features associated with host gene regulation. Our results illustrate that single-cell analyses of viral infection, even in cell culture, provide meaningful insights into host-pathogen interactions during viral infection.

## METHODS

### Cell culture of MRC-5

MRC-5 human lung fibroblast cells (ATCC, #CCL-171) were thawed, passaged, expanded, and seeded for viral infection experiments using E-10% Culture Medium containing: Eagle’s Minimum Essential Medium (EMEM) containing 2 mM L-Glutamine, 1 mM sodium pyruvate, and 1500 mg/L sodium bicarbonate (ATCC, #30-2003), supplemented with 10% (v/v) filter-sterilized Fetal Bovine Serum (FBS: ATCC, #30-2020). MRC-5 cells were incubated in water-jacketed cell culture incubators at 37 °C, 95% relative humidity, and supplied with 5% CO_2_. Several aliquots of 10^6^ MRC-5 cells were cryopreserved for infection experiments at passage #2 from the original source vial using ice-cold EMEM supplemented with 25% (v/v) filter-sterilized FBS and 5% (v/v) Dimethyl Sulfoxide (DMSO: ATCC, #4-X).

#### Virus batch propagation and validation

Human coronavirus strain 229E (HCoV-229E) from ATCC ((#VR-740, Lot #70053995) was used for in-house virus batch propagation. Virus inoculum was made by diluting HCoV-229E viral stock in *E-2% Infection Medium* consisting of EMEM supplemented with 2% (v/v) filter-sterilized FBS. MRC-5 cells were seeded into 175 cm^2^ tissue-culture flasks using 5.0 x10^6^ cells per flask (2.8 x10^4^ cells/cm^2^). MRC-5 cell monolayers were infected at MOI=1 using 12 mL virus inoculum for 1 hr at 35 °C, then 17 mL *E-2% Infection Medium* was added prior to further incubation at 35 °C. After 3 days, when severe cytopathic effects were observed in a majority of cells, the virus-containing supernatant was collected from each flask (∼100mL total), centrifuged at 4000 rcf for 10 min at 4 °C to remove cell debris, then filtered through a 0.22 um PES membrane filter (Millipore, #S2GPT05RE). The virus batch was stored long-term as 5 mL aliquots at −80 °C. The in-house HCoV-229E virus batch was validated by titration and infection of MRC-5 cells. Cell viability was assayed by microscopy for 3 days post-infection using the reagents and instructions provided in the Live/Dead Cell Staining kit (Invitrogen, #L3224).

### Large-scale viral infection and cell harvesting

HCoV-229E virus inoculum was prepared in *E-2% Infection Medium* using frozen virus stock aliquots immediately before infection. After 6 days of incubation, seeded flasks of MRC-5 cells were infected at MOI=4 using 13 mL virus inoculum for 1 hr at 35 °C, and then 15 mL *E-2% Infection Medium* was added prior to incubation at 35 °C for 24-48 hr. Mock-infected flasks were given 13 mL *E-2% Infection Medium* without virus for 1 hr, then another 15 mL *E-2% Infection Medium* before incubation in parallel with the virus-infected flasks for 24-48 hr.

A total of 36 flasks of MRC-5 cells were divided among treatments, mock-infected (12 flasks) and virus-infected (24 flasks), and then into batches with associated time points: A-24 hr, A-48 hr, B-24 hr, B-48 hr, C-24 hr, C-48 hr, each with 2 flasks for mock-infected and four flasks for virus-infected. Infections were staggered by 3 hr each to enable accurate timing for cell harvests at 24 hr and 48 hr time points. Mock-infected and virus-infected cells were harvested one batch at each time point: A-24, then B-24 3 hours later, then C-24 3 hours later. 24 hours after the first batch harvest (A-24), A-48 was harvested, then B-48 3 hours later, then C-48 3 hours later. C batch harvesting was interrupted resulting in a loss of the harvested cells (12 flasks). For each batch at each time-point, cells were washed once using 1X PBS, detached for 25 min at 37 °C using StemPro Accutase, and thoroughly resuspended in room-temperature *E-10% Culture Medium* before cell counting with the Countess instrument. Mock-infected and virus-infected cell suspensions were aggregated among biological replicates (2 flasks for Mock, 4 flasks for Viral Infection) and then divided into technical replicates and cryopreserved in ice-cold *Freeze Medium* for downstream ATAC-seq and scRNA-seq techniques. All data presented is from analysis of biological replicate flasks from A-24, B-24 and A-48, B-48.

### Library preparation: scRNA-seq and ATAC-seq

For single-cell RNA-seq (scRNA-seq), triplicate aliquots of 2×10^5^ mock-infected and virus-infected cells from two biological replicates (A and B) under each condition (Mock and 229E Infected), for both timepoints 24 and 48 hpi, were thawed ice-cold *E-10% Culture Medium* containing 1X HALT Protease Inhibitor Cocktail (Invitrogen, #78430). Cells were counted to achieve 1×10^5^ viable cells per sample, then cross-linked, permeabilized, and cryopreserved using the reagents and instructions provided in the Evercode Fixation Kit (Parse Biosciences, #ECF2101). Cross-linked cells were processed using the Evercode WT Mini v2 Kit (Parse Biosciences, #ECW02110). Cross-linked cells were subjected to four rounds of barcoding and labelled using combinatorial indexing. Sub-libraries containing 25K and 10K cells each were template-switched, amplified, and cleaned using *SPRI Beads.* Libraries underwent fragmentation, end-repair, A-tailing, and sub-library indexing PCR was performed using the *Unique Dual Index (UDI) Plate*. Finally, libraries underwent post-amplification double-sided SPRI size-selection. The concentration and size of the libraries were verified using the TapeStation DNA 5000 Assay Reagents and Screen Tapes (Agilent, #5067-5589 and #5067-5588). Sub-libraries were sequenced on NextSeq 500 HO flow cell to generate paired-end reads where read 1 is sequenced for 221 cycles and read 2 for 86 cycles using the using the NextSeq 500 High Output Kit v2.5 (300 cycles) (Illumina, #20024908) at the Los Alamos National Laboratory Genome Center.

For ATAC-seq, triplicate aliquots paired to the scRNA-seq samples, each for A-24 Mock, B-24 228E, A-48 Mock, and B-48 229E, were thawed into ice-cold *E-10% Culture Medium* containing 1X HALT Protease Inhibitor Cocktail (Invitrogen, #78430). Cells were counted to achieve 70,000 viable cells per sample, then processed for ATAC-seq analysis using the Active Motif ATAC-seq Kit (#53150) as previously described^20^. Briefly, cells were washed in ice-cold 1X PBS, then lysed in ice-cold *ATAC Lysis Buffer* (from kit), and centrifuged for 10 min at 4°C. Cell lysates were incubated with *Tagmentation Master Mix* (from kit) to prepare tagmented DNA, which was then purified using DNA purification columns (from kit). DNA libraries were prepared from tagmented DNA amplified via PCR using Q5 Polymerase and unique combinations of i7 and i5 Indexed Primers, PCR amplified, and cleaned using magnetic DNA purification beads. Bead-based size selection was performed using AMPureXP Reagent to retain fragments between 150 and 600 bp. DNA libraries were quantified using a high-sensitivity Qubit dsDNA Quantification Assay kit (Thermo Fisher Scientific, #Q32851) and KAPA Library Quantification Kit (Roche, #KR0405), then sequenced via the Illumina NextSeq 2000 sequencer platform in paired-end mode at the Los Alamos National Laboratory Genome Center.

### scRNA-seq bioinformatic analyses

#### Pre-processing

Single-cell RNA-seq libraries were sequenced in multiple runs and reads were concatenated in a sublibrary specific manner. Reads were mapped to a concatenated human (T2T) and human coronavirus 229E (NCBI: NC_002645.1) genome using ParseBiosciences-Pipeline.1.1.2. The number of viral reads per sample are presented in Table S1. The resulting unfiltered count matrix was processed using CellBender to identify barcodes representing real cells and remove background noise^21^. We trained the model for 50 epochs, and set the number of expected cells to 25,000, the total number of droplets included to 40,000, and the FPR to 0.05. The resulting filtered count matrix was then processed further using the Scanpy v1.10.3^22^. Only cells with a cell probability of ≥ 0.9999 and reads between 5,000 and 30,000 were kept in the analysis (Figure S1). 99.6% of infected cells contained mapped viral reads. Post QC, we analysed profiles of 13,913 cells across all samples, including 4,192 mock-infected and 2,724 virus-infected cells at 24 hpi and 3,593 mock-infected and 3,404 virus-infected cells at 48 hpi. We performed read-depth normalization using the scanpy.pp.normalize_total() function and performed a log(X + 1) transformation using the scanpy.pp.log1p() function. Next, we selected the top 5000 most variable genes with scanpy.pp.highly_variable_genes() using the “Seurat V3” flavor. Next, we computed the PCA decomposition and the nearest neighbour graph, constructed a UMAP-embedding, and performed Leiden clustering. To annotate cell cycle stages, we ran <u>sc.tl</u>.score_genes_cell_cycle() using the cc.genes.updated.2019$g2m.genes and cc.genes.updated.2019$s.genes in seurat v5^23^.

#### Responsive gene identification

To identify genes that respond to viral infection, we performed Wilcox rank-sum differential expression testing across several contrasts. To identify viral exposure-related genes, we performed differential expression testing using sc.tl.rank_genes_groups() function across several contrasts, namely 229E versus Mock at 24 hpi, 229E versus Mock at 48 hpi, and 229E 24 versus 48 hpi. To identify genes related to heterogeneity within infection conditions, we performed cluster-level differential expression testing within the Mock control clusters (Mock.1, Mock.2, Mock.3, Mock.4, Mock.5, Mock.6, Mock.7), within the 24 hpi clusters (24hpi.1, 24hpi.2, 24hpi.3), and within the 48hpi clusters (48hpi.1, 48hpi.2). Furthermore, to identify genes associated with viral load in infected samples, we performed a likelihood ratio test (LRT) in DESeq2^24^, comparing the full model ∼ replicate + timepoint + viral_load to the reduced model ∼ replicate + timepoint, thereby testing the contribution of viral load after accounting for replicate and timepoint effects. To identify changes in cell-cell communication between treatment conditions, cell clusters, and low-viral-load (putative bystander-like) and higher-viral-load (infected) cells, we used NICHES^25^ with the FANTOM5 reference database, following their Differential CellToCell Signaling Across Conditions vignette (https://msraredon.github.io/NICHES/articles/04%20Differential%20CellToCell.html).

#### Predictive expression networks

In addition, we constructed single-cell PENs using scRNA-seq data from mock controls, 24hpi, and 48 hpi using iRF-LOOP^26^. To prepare the expression data for scPEN construction, we remove low variance genes and select a single representative gene for groups of genes that have Pearson’s correlation values > 0.95. Next, we run iRF-LOOP to construct the scPEN networks. In post-processing, we removed any edges derived from models with an R^2^ < 0.5 and kept the top 5% of edges based on edge weight. To describe the total change in a gene’s connectivity upon viral exposure, we computed a rewiring score, RS, defined as the sum of the absolute differences in weights across all edges for a given gene. In addition, we compute the degree centrality, eigenvector centrality, and betweenness centrality for each scPEN using the degree_centrality(), eigenvector_centrality(), and betweenness_centrality() functions from networkx^27^. To run gene set enrichment analyses for our DEG gene sets based on the rewiring and centrality-based rankings, we use the gseapy.prerank() function. All functional enrichment analyses were performed with gprofiler-official^28,29^ using only detected genes as a background.

#### ATAC-seq bioinformatics analyses

Twelve ATAC-seq libraries were sequenced as described above and data was analyzed as previously described^20,30^. Briefly, raw reads were trimmed to remove Nextera adaptors. Reads with repetitive sequences were filtered using Fastp^31^ and bases with low quality scores (q < 15) were also removed. Processed reads were aligned to the telomere-to-telomere (T2T) human reference genome, version 2^32^ using BWA^33^. Duplicate sequenced pairs and mitochondrial reads were marked and removed from analysis using Samblaster^34^. Samples were further filtered using samtools with the following flags: -F 4 -F 256 -F 512 -F 1024 -F 2048 -q 30. Loci displaying significant enrichment of paired-end reads were identified as peaks using MACS2^35,36^ while processing reads in BAMPE mode. Replicate correlation was assessed using deepTools^37^ - plotCorrelation function on SPMR normalized bigwig files. TSS-centered accessibility profiles were generated using the computeMatrix function from deepTools. Open chromatin regions (OCRs) were defined by merging MACS2-called peaks using BEDtools^38^. Computation of the PCA decomposition of raw reads across consensus peaks and identification of differentially accessible OCRs using identification for matrix design (∼time + infection) was performed using DESeq2^24^. Raw reads from all individual samples across consensus peaks (i.e., merged intervals of MACS called peaks) per feature were used as input for PCA. To determine the global levels of feature profile distribution in control and treated samples, –computeMatrix function from deepTools with –binsize 10 kb was used. SPMR-normalized read coverage around midpoints of corresponding consensus peaks (i.e., merged intervals of MACS called peaks) were estimated. TFBS enrichment was performed using Homer using only known vertebrate motifs^39^.

#### Response gene characterization with MENTOR

Given the differential expression statistics, a subset of genes was selected to partition using MENTOR^40^, a tool for functional partitioning of genes based on multiple lines of interaction evidence embedded in a multiplex. For each contrast, we select the most extreme |log_2_FC| value in any of the contrasts and select all genes with a |log_2_FC| value > 7.5. MENTOR software was run using a multiplex consisting of the HumanNet V3 CX, PD, GI, and GN networks^41^, a HumanNet V3 + STRING protein-protein interaction network^42^, a human lung predictive expression network (PEN) constructed from GTEx data^43^, and 2 human lung fibroblast single-cell PENs constructed from GTEx data^44^. We applied MENTOR to obtain clades of functionally connected genes for further interpretation.

## RESULTS

### Single-cell RNA-seq analysis reveals the heterogeneity of viral progression in infected MRC-5 cells

To investigate the heterogeneity of host responses, we performed large-scale infections of MRC-5 lung fibroblasts with human coronavirus 229E (HCoV-229E), and profiled gene expression using single-cell RNA sequencing and chromatin accessibility using ATAC-seq at 24 and 48 hours post-infection (hpi) (Figure 1A). We determined the viral load, defined as the fraction of transcripts mapping to the viral genome, as an indication of viral infection. Contrary to our expectations, infected cells at 24 hpi had a higher viral load than cells at 48 hpi (Figure 1B). This finding suggests potential survivorship bias, i.e., cells that survive at 48 hpi either did not experience the infection or successfully eliminated it. Additionally, 2.9% of cells at 24 hpi and 3.1% of cells at 48 hpi contained fewer than 100 viral transcripts (Figure S2). These cells, in which macromolecule biosynthetic process genes and unfolded protein response genes exhibit lower expression than in infected cells (Table S2, S3), were either never infected or were particularly resilient to infection. The distribution of reads mapping to HCoV-229E across infected cells revealed heterogeneity in viral transcription rate or a lack of uniformity in viral proliferation (Figure 1B).

**Figure 1.**
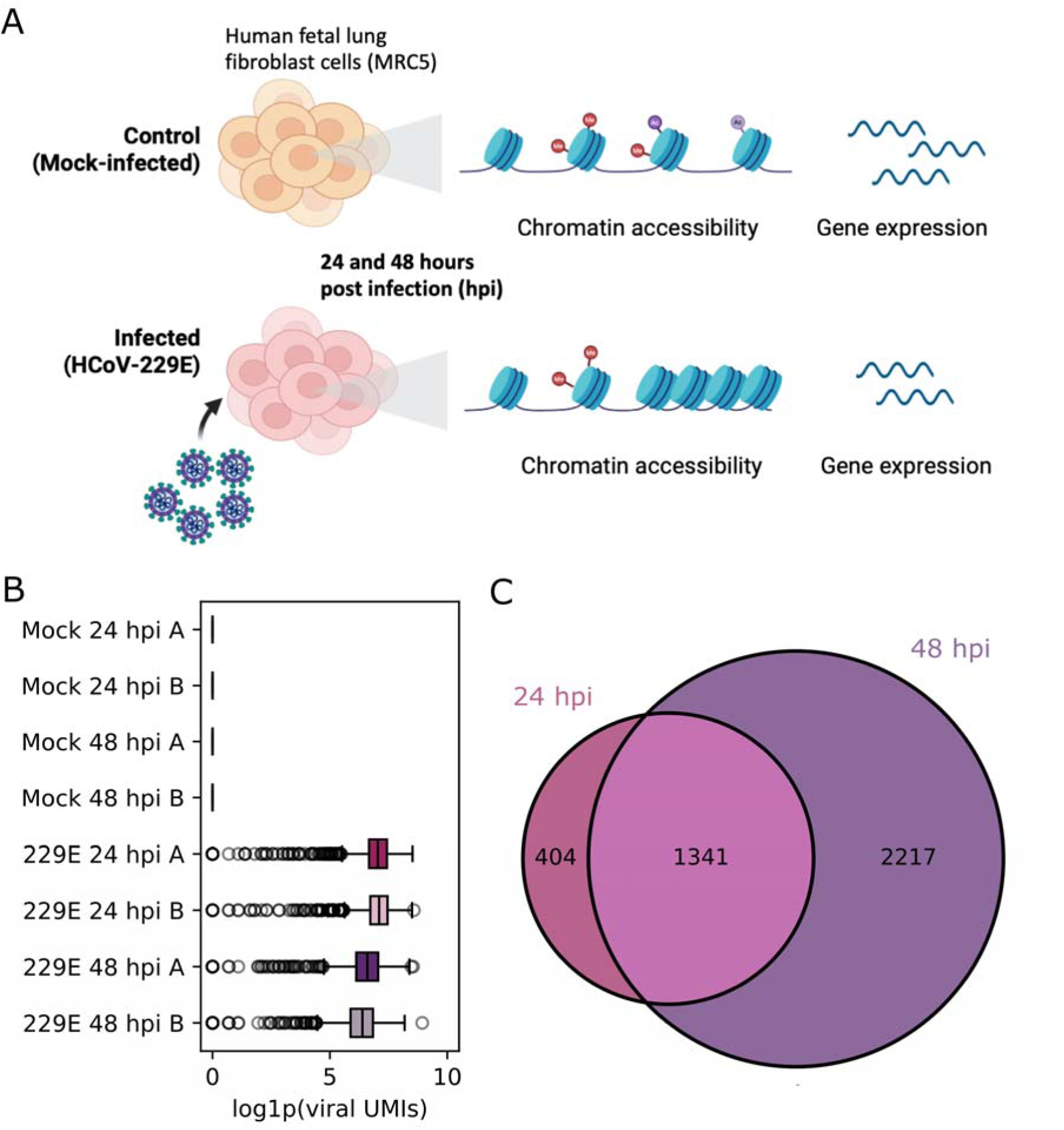
Analysis of Coronavirus 229E infection of MRC-5 cells. **A)** Experimental protocol with sample collection at 24 and 48 hours post infection (hpi) for two duplicate experiments (A and B). For scRNA-seq, this resulted in transcriptomes from 13,913 cells across all samples, including 4,192 mock-infected and 2,724 virus-infected cells at 24 hpi and 3,593 mock-infected and 3,404 virus-infected cells at 48 hpi. **B)** The viral load across samples after quality control. The median log1p(viral reads) detected in 24 hpi sample A is 7.0 (SD 1.0), in 24 hpi sample B is 7.1 (SD 0.9), and across samples A and B is 7.1 (SD 1.0). The median log1p(viral reads) detected in 48 hpi sample A is 6.6 (SD 1.0), in 48 hpi sample B is 6.4 (SD 1.0), and across samples A and B is 6.5 (SD 1.0). **C)** Venn diagram representing the number of differentially expressed genes in a cell-level, time-matched 229E-infected versus mock-control differential expression test at a threshold of adjusted p-value < 0.001 and |log_₂_ FC| > 2.

Although the average number of viral reads was lower at 48 hpi than 24 hpi, we identified more differentially expressed genes (DEGs) at 48 hpi from the time-matched individual cell-level differential expression analysis (Figure 1C, Table S2). At both timepoints, upregulated genes were enriched for transcriptional and biosynthetic regulation, stress response and proteostasis, immune and inflammatory signaling, apoptosis, and signaling and cell communication (Table S3). In contrast, downregulated genes were enriched for collagen metabolic process, extracellular matrix (ECM), development and differentiation, signaling, cytoskeleton and motility, and adhesion illustrating the virus-induced shift from fibroblast-like homeostatic functions towards a stress-adaptive, antiviral state in the MRC-5 cells. A subset of heat shock, interferon-mediated antiviral defense, and stress- and metabolism-related adaptation genes showed exceptionally high log_2_ fold-changes (log_2_FC > 20) at 24 and 48 hpi. Similarly, by 48 hpi, extreme upregulation was also observed for several genes involved in NF-κB-mediated inflammation, extracellular matrix organization, cell adhesion, and tissue remodeling, despite lower viral load. This response was paired with significant downregulation (log_2_FC < −20) of select genes with known functions in calcium signaling, cell adhesion, and vesicle transport at 48 hpi. Taken together, these results illustrate that HCoV-229E infection induced broad transcriptional reprogramming of MRC-5 cells, characterized by a transition from a fibroblast-like, matrix-producing state toward a stress-adapted, metabolically reprogrammed, antiviral state; this reprogramming was predicated on viral load variability among cells in culture.

### Heterogeneity of the transcriptomic response

To further investigate the heterogeneity in the transcriptomic host response to infection, we performed clustering analysis of the scRNA-seq data (Figure 2A). In addition to differences in viral read distributions between samples (Figure 1B), viral read distributions also differ across transcriptomically defined clusters, particularly at 48 hpi (Figure 2B). The relatively balanced contribution of cells from each sample to each of these clusters illustrates that cluster-level viral load differences were not driven solely by sample composition (Figure S3) but may instead reflect distinct stages of viral progression within individual cells. At 24 hpi, cells partition into three clusters (24hpi.1, 24hpi.2, and 24hpi.3). Cluster 24hpi.2 expressed a marginally higher number of viral reads, with upregulated genes enriched for the unfolded protein response and downregulated genes enriched for cell cycle and proliferation control (Table S3). While clusters 24hpi.1 and 24hpi.3 had similar viral load distributions (Figure 1B), they had transcriptional differences, where downregulated genes in cluster 24hpi.1 and upregulated genes in cluster 24hpi.3 were enriched for cell cycle processes (Table S3). At 48 hpi, cells partition into two clusters (48hpi.1 and 48hpi.2). Cluster 48hpi.1 had a higher viral read percentage, and upregulated genes were enriched for the unfolded protein response, whereas upregulated genes in cluster 48hpi.2 were enriched for cell cycle-related terms (Figure 2B). In concordance with these results, a likelihood-ratio test revealed that genes that are significantly positively associated with viral load are enriched for unfolded protein response, apoptosis, and metabolic adaptation, while genes that are negatively associated with viral load are enriched for cell cycle, cytoskeletal processes, cell migration, and glycolysis (Table S4, Table S5). However, computed cell cycle scores were only weakly negatively correlated with viral load (ρ = −0.15 for S-scores, and ρ = - 0.12 for G_2_/M-scores).

**Figure 2.**
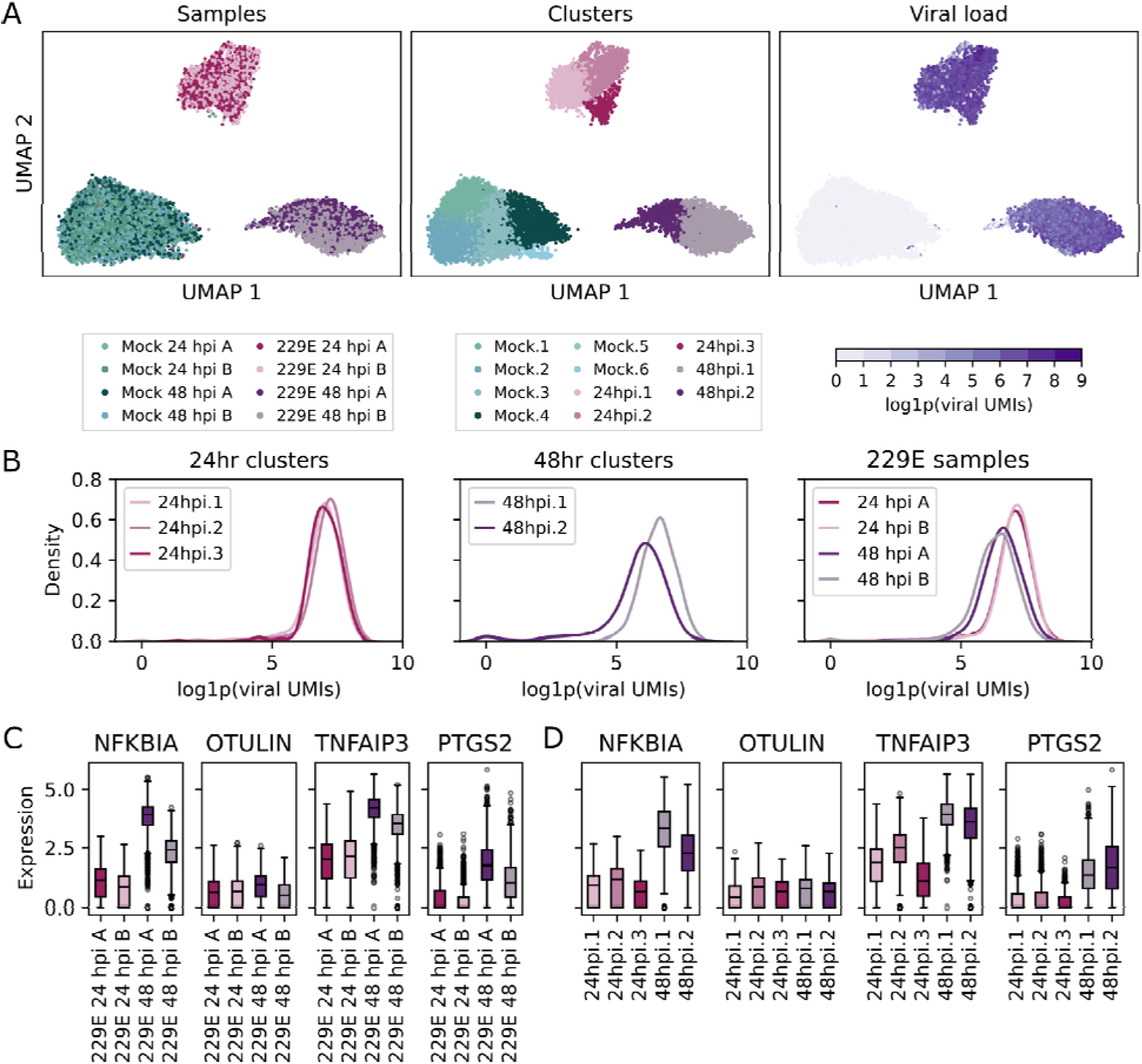
Cellular heterogeneity of the transcriptomic response to coronavirus 229E infection. **A)** UMAP plots colored by sample (left), cluster (middle), and viral load (right) for all samples in all clusters. **B)** Viral read distributions in clusters at 24 hpi (left), clusters at 48 hpi (middle), and per sample (right). At 24 hpi, the median log1p(viral reads) was 7.2 (SD 0.8) in cluster 24hpi.2, 7.0 (SD 0.9) in 24hpi.1, and 7.0 (SD 0.7) in cluster 24hpi.3. At 48 hpi, the viral read percentage was 6.6 (SD 0.7) in cluster 48hpi.1, and 6.0 (SD 1.3) in cluster 48hpi.2. The median log1p(viral reads) was 7.0 (SD 1.0) in sample A, and 7.1 (SD 0.9) in sample B. At 48 hpi, the median log1p(viral reads) was 6.6 (SD 1.0) in sample A, and 6.4 (SD 1.0) in sample B. **C)** Expression levels of key NF-κB regulators and effectors across virus-infected samples. **D)** Expression levels of key NF-κB regulators and effectors across virus-infected clusters.

In the mock-control condition, transcriptomic heterogeneity primarily reflected cell cycle state. Clusters Mock.4 and Mock.6 overexpressed genes associated with active cell cycling compared to clusters Mock.1, Mock.2, Mock.3, and Mock.4 (Table S3, Figure S4). Differences in the proportion of cycling cells between biological replicates were minimal (Figure S5). Infected samples contained fewer cycling cells than mock-control samples, and this proportion decreased slightly from 24 to 48 hpi, whereas mock-control samples show a modest increase in cycling cells at 48 hpi. This reduction in cycling was concurrent with the increased expression of G_1_-markers *CDKN1A* and *CDKN1B* suggesting G_1_ arrest (Figure S4). Together, these results imply that while cell cycle state contributes to transcriptional heterogeneity, particularly in the mock-control condition, it does not fully explain the divergent responses observed among infected cells. Instead, infection-induced responses, specifically the upregulation of the unfolded protein response, was potentially a defining source of transcriptional variation as viral replication progressed.

We also observed variability in viral read percentages among biological replicates from the same time points (Figure 2B). To investigate the extent to which these sample-level variations impacted the transcriptomic response to infection, we performed differential expression testing between samples at 24 and 48 hpi (Table S6). Though there were very few DEGs, the NF-κB inhibitor *NFKBIA* was upregulated in 48 hpi sample A compared to sample B, along with less pronounced increases of several NF-κB regulators (*OTULIN*, *TNFAIP3)* or effectors of NF-κB (*PTGS2*) (Figure 2C). Beyond these sample-specific expression patterns, the negative regulators of NF-κB, *NFKBIA* and *TNFAIP3*, were overexpressed in the high-viral-load clusters 24hpi.2 and 48hpi.1 (Figure 2D). Both of these genes also show higher expression in low-viral-load cluster 24hpi.1 (cell cycle-depleted) than in cluster 24hpi.3, which also has a low viral load, but is enriched in cell cycle genes. These results suggest that inhibition of NF-κB increases when viral load is high (regardless of the timing in the culture) or when cell proliferation capacity is reduced. This pattern underscores NF-κB’s central, context-dependent role in shaping the variability of HCoV-229E infection rates and the associated transcriptional responses observed both within cultures and between biological replicates.

### HCoV229E infection induces changes in cell-to-cell communication

Differential expression analysis revealed variation across infection conditions, samples, and clusters in individual cells that impact structural integration, immune modulation, and stress responses. As many of these processes involve cell-to-cell signaling, we compared known human ligand-receptor interactions between clusters and samples using NICHES (see Methods). In high-viral-load clusters, at 24 hpi, NICHES-inferred interactions between ECM remodeling and integrin-associated genes were downregulated while interactions between growth factor, inflammatory, and cell-cell signaling genes increased compared to lower viral-load clusters (Table S7). Between mock and infected cells and from 24 to 48 hpi, NICHES-inferred interaction activity showed a consistent decrease in ECM- and adhesion-related interactions and an increase in growth factor and inflammatory signaling-mediated interactions. In addition, we observed a reduction in IL11-IL6ST interactions, consistent with decreased IL-11-associated gp130 signaling, along with reduced ECM-integrin-related and tissue homeostasis-related interactions. These findings illustrate that infection shifts cell-cell communication from structural maintenance toward immune activation and compensatory repair mechanisms within *in vitro* cell cultures.

### HCoV-229E infection rewires gene networks

For data-driven identification of putative changes in gene-gene interactions upon viral exposure, we compared single-cell predictive expression networks (scPENs) from mock control samples, 229E 24 hpi samples, and 229E 48 hpi samples independently. To evaluate the extent to which a gene’s network connections were reorganized between conditions, we computed a rewiring score, which quantifies the extent to which a gene’s interactions change upon viral infection at 24 hpi, 48 hpi, and between 24 and 48 hpi (see Methods, Table S8). Across comparisons, cell cycle genes were found to be most strongly rewired, suggesting that the infection-induced G_1_-arrest identified in differential expression analyses is caused by coordinated changes in expression of cell cycle-related genes. Additionally, gene set enrichment analysis (GSEA) showed strong enrichment for viral-load associated genes (ES = 0.91 - 0.93) and upregulated genes (ES = 0.89). Notably, NF-κB regulators (*NFKBIA*, *NFKBIZ*, *TNFAIP3*, and *TRAPPC9*) were among the most highly rewired genes at 48 hpi and between 24 and 48 hpi (Table S8). We also investigated the correlation between rewiring scores with degree, eigenvector and betweenness centrality, where degree centrality measures how many connections each gene has, eigenvector centrality captures the extent to which a gene is connected to other highly connected genes, and betweenness centrality quantifies how often a gene lies on the shortest paths between other genes, reflecting its role as a bridge within the network. Rewiring scores were also strongly correlated with degree, eigenvector, and betweenness centrality in mock networks (ρ = 0.70–0.82), and hub genes were significantly more rewired than non-hubs (p < 0.001). These results illustrate that rewiring preferentially affects central nodes, and that these genes show virus-induced expression changes.

Though rewiring scores correlated with centrality in mock control networks, rewiring was not driven by increased connectivity. Namely, rewiring scores were negative or weakly correlated with changes in degree (ρ = −0.54 to −0.33) and eigenvector centrality (ρ = −0.27 to −0.13) and showed minimal association with changes in betweenness centrality (ρ < 0.15). This suggests that gene-gene connections weaken or are redistributed rather than gained upon viral exposure. However, though global correlations between rewiring scores and changes in centrality were weak, select highly rewired genes, including *ABL2,* a critical host factor required for efficient SARS-CoV and MERS-CoV replication^45^, exhibited strong changes in eigenvector centrality, highlighting gene-specific coupling between local rewiring and global topology. Functionally, genes losing degree centrality were enriched for cell cycle processes while those gaining degree centrality were enriched for negative regulation of mitotic cell cycle and checkpoint signaling at 24 hpi and stress related and inflammatory signaling pathways like the unfolded protein response, glycolysis, and IL-17/TNF signaling by 48 hpi. Similarly, genes with increased eigenvector centrality at 24 hpi and those with decreased eigenvector centrality at 48 hpi were enriched for cell cycle pathways, while genes with increased eigenvector centrality at 48 hpi were enriched for stress-related stress related and inflammatory signaling pathways. Finally, genes with decreased betweenness centrality at 24 and 48 hours were enriched for cell cycle processes, but genes with increased betweenness centrality from 24 to 48 hours were enriched for inflammatory signaling pathways. Genes with negative changes in betweenness centrality at 24 hpi were enriched for cytoskeletal architecture, Rho GTPase signaling, and chromosome organization and for cell cycle at 48 hpi. In addition, positive viral load associated genes preferentially gained centrality, while negatively associated genes preferentially lost centrality. Together, these results indicate that infection-induced rewiring primarily affects network hubs, and induces a shift from cell cycle-related host signaling to stress response signaling through coordinated redistribution of connectivity.

### Infection-induced changes of the MRC-5 chromatin landscape

Given our observations that viral proteins may regulate genes involved in chromatin remodeling, we investigated the concordance between changes in transcription and chromatin accessibility. We performed bulk ATAC-seq profiling of HCoV-229E-infected MRC-5 cells and their time-matched mock controls at 24 and 48 hpi (Figure 1A). Median accessibility around transcription start sites (TSSs) decreased upon viral infection at 24 hpi, with less pronounced differences at 48 hpi (Figure 3A). The difference was more pronounced when considering the mean accessibility profile around the TSS (Figure 3B) than the median profile, suggesting that a subset of regions becoming inaccessible drove the observed overall decrease in TSS-proximal accessibility. Similarly, while at 48 hpi median accessibility further from the TSS increased marginally upon infection, mean accessibility did not, suggesting that there was a subset of inaccessible distal regions masking an overall increase in distal chromatin accessibility.

**Figure 3.**
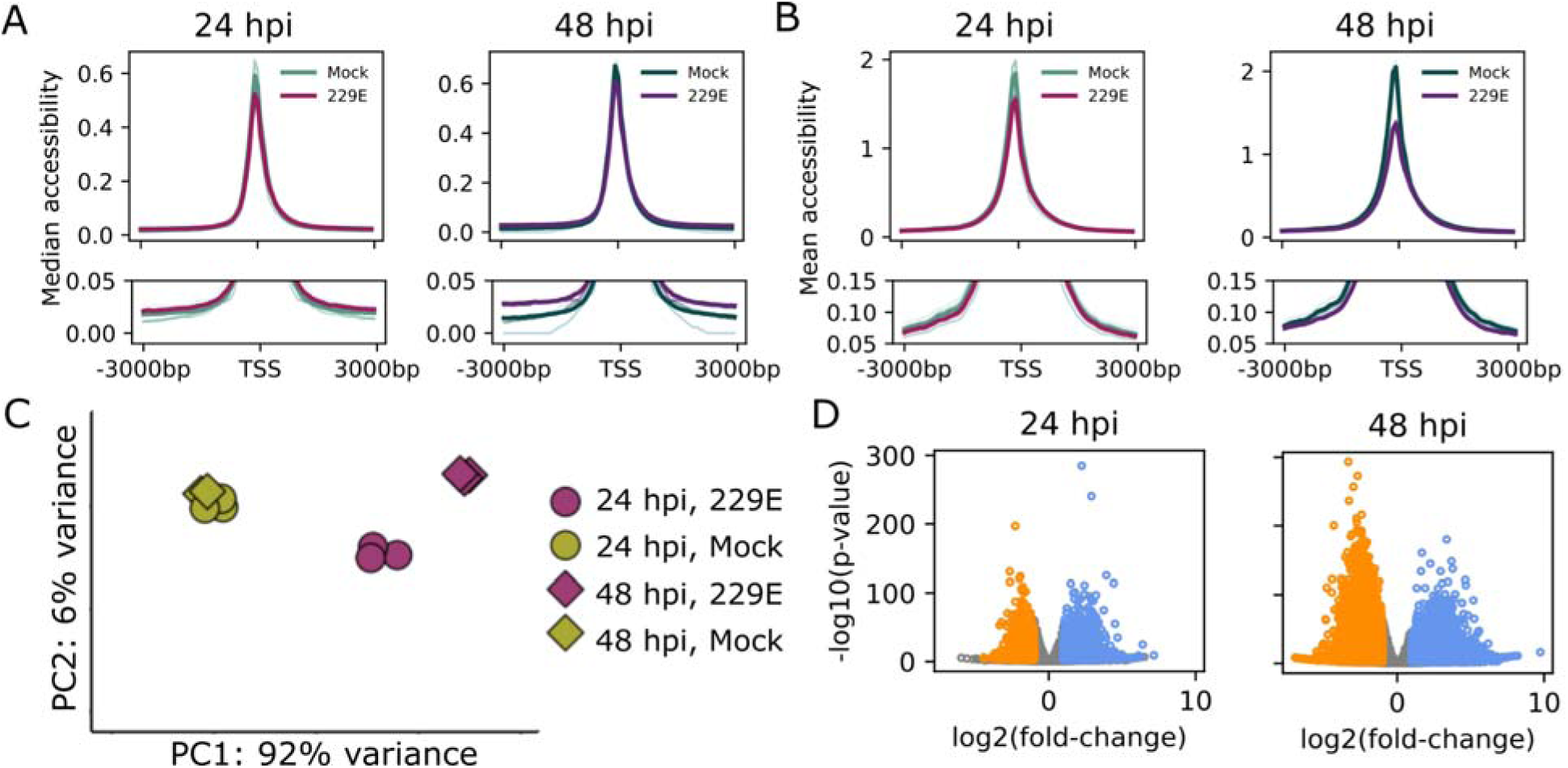
HCoV-229E impacts on bulk MRC-5 chromatin accessibility profiling at 24 and 48 hpi. **A)** The median aggregated accessibility profiles centered on the transcription start sites (TSS; ±3kb) for mock and 229E-infected samples at 24 and 48 hpi. The bold lines show median and mean profiles computed over mean profiles across all samples, while lower opacity profiles show variability across replicates. **B)** The mean aggregated accessibility profiles centered on the transcription start sites (TSS; ±3kb) for mock and 229E-infected samples at 24 and 48 hpi. The bold lines show median and mean profiles computed over mean profiles across all samples, while lower opacity profiles show variability across replicates. **C)** Principal component analysis (PCA) of accessibility profiles separates mock-control and 229E infected samples at 24 and 48 hpi. Axes indicate the variance explained by each principal component, marker shapes indicate time point, and marker colors indicate infection status. **D)** Volcano plots of differential accessibility (infected versus mock-control) at 24 (left) and 48 (right) hpi. Each point is an open chromatin region, and blue points indicate significantly increased accessibility, while orange points indicate significantly decreased accessibility. Gray points are not significant. At 24 hpi, 6% of open chromatin regions (OCRs) show increased chromatin accessibility (blue) and 5% show decreased chromatin accessibility (orange). At 48 hpi, 20% of OCRs show increased chromatin accessibility (blue), and 13% show decreased chromatin accessibility (orange).

There was a marked time-dependent effect of HCoV-229E infection on global chromatin accessibility (Figure 3C) driven by bidirectional changes in chromatin accessibility at both 24 and 48 hpi (Figure 3D). Approximately 6% of differentially accessible regions between infected cells and mock controls at each time point overlapped gene promoters (defined as −1000 to +100 bp from the TSS). Notably, 66.8% of differentially accessible promoters at 24 hpi retain their differential accessibility at 48 hpi. Next, we evaluated the relationship between promoter accessibility and differential gene expression. While only 4.0-5.3% of genes with differentially accessible promoters are differentially expressed, the majority of these genes (57.1-88.5%) follow the expected directionality of differential expression, i.e., higher accessibility is associated with greater gene expression (Table S9). Similarly, when we examine extremely upregulated genes (log_2_FC > 20), we find that a substantial proportion (12.8% at 24 hpi, and 19.3% at 48 hpi) have differentially accessible promoters). Overall, these results indicate that while changes in promoter accessibility do not perfectly mirror gene expression directionality, a broad concordance between the two is evident during infection.

To explore potential functional implications of differential chromatin accessibility, we first performed overrepresentation analysis on genes with differentially accessible promoters. We find enrichment of RNA processing and IL-17 signaling for promoters that become accessible at 24 hpi, followed by enrichment of membrane transport at 48 hpi, alongside enrichment of REST/NRSF-transcription factor binding motifs in genes with promoters that become inaccessible at 24 and 48 hpi (Table S10). In addition, we performed transcription factor binding site (TFBS) enrichment analysis (Table S11). At 24 hpi, regions that became accessible were enriched for NF-κB subunits, AP-1/ATF/CEBP family stress-response transcription factors, IRF-mediated immune regulators, TEAD factors associated with Hippo signaling, circadian regulators, and a broad set of developmental and homeobox transcription factors. At 48 hpi, enrichment for NF-κB subunits and stress response TFs persisted, accompanied by homeobox and an insulin regulator Pdx1, suggesting additional engagement of metabolic and morphogenesis programs. Importantly, the sustained enrichment of NF-κB-related genes and their accessibility reinforces the single-cell transcriptomics-informed hypothesis that NF-κB signaling plays a central role in regulating the response to HCoV-229E in MRC-5 cells predicated on viral load.

### MENTOR identifies host response signaling to HCoV-229E infection

We selected a stringent set of differentially expressed genes for further exploration using MENTOR^15^, a tool we previously developed for identifying functional gene modules based on a multiplex network embedding known human biology (Figure 4, see Methods). The accompanying heatmap illustrates the log2FC values for the considered differential expression contrasts (Figure 4, tracks I-IV) and the direction of the significant infection-induced differential accessibility of gene promoters (Figure 4, track V). The resulting functional partitioning revealed that most modules consist of genes with strong differential expression due to infection (M1-12, M14-15, M18-20); these modules were enriched for pathways involved in lipid processing (M2), transmembrane transport (M2, M3, M5), antimicrobial response and chemokine signaling (M6), cytokine signaling (M4, M6, M18), NF-κB signaling and broader immune responses (M18), cytoskeletal remodeling (M11), and HSF1-mediated heat shock response (M15) (Figure 4, Table S12). In several of the modules– transport module M2, chemokine/cytokine and antimicrobial response module M6, cytoskeleton module M11, and HSF1-mediated heat shock response module M15–at least 25% of genes had differentially accessible promoters (Figure 4, track V). While we do not observe chromatin accessibility changes in the promoters of all differentially expressed genes, MENTOR findings support the perspective that transcriptional response to infection is, at least in part, coordinated through some level of regulation of chromatin accessibility, particularly for genes with substantial changes in gene expression (Figure 3C). One of the remaining modules, M16, showed more differential expression in mock-control cell clusters accompanied by enrichment of cell cycle-related processes (Figure 4, Table S12). Notably, several genes in these cell cycle-dominated clades were upregulated in the lower viral load clusters 24hpi.3 and 48hpi.2, further substantiating that viral infection interferes with the proliferative capacity of MRC-5 cells. Finally, the two remaining modules, M13 and M17, represent mechanistic bridges between these processes and contain genes originating from both infection and mock-control cell clusters. More specifically, module M13 was enriched for tubulin binding, microtubule organization, ATP/adenyl nucleotide binding, and polymeric cytoskeletal fibers, suggesting a link between viral infection and cell cycle–associated cytoskeletal dynamics. Module M17 was enriched for nucleosome and chromatin organization, histone modification pathways, and DNA replication/repair processes, suggesting a link between epigenetic regulation of cell function and chromatin remodeling during viral infection. Additionally, module M17 includes *TCEAL7*, a negative NF-κB regulator that promotes transcription of cyclin D, a key G_1_/S transition regulator. Furthermore, M8 shows that the previously mentioned NF-κB modulators *NFKBIA* and *TNFAIP3* are upregulated in Mock.2 in addition to infected cells. This *a priori* knowledge-centric gene interpretation identified NF-κB as a central player in the mechanistic link between inflammatory responses and cell cycle regulation in response to viral infection, providing further evidence for the key role of NF-κB in governing the heterogeneous response of MRC-5 cells to HCoV-229E infection.

**Figure 4.**
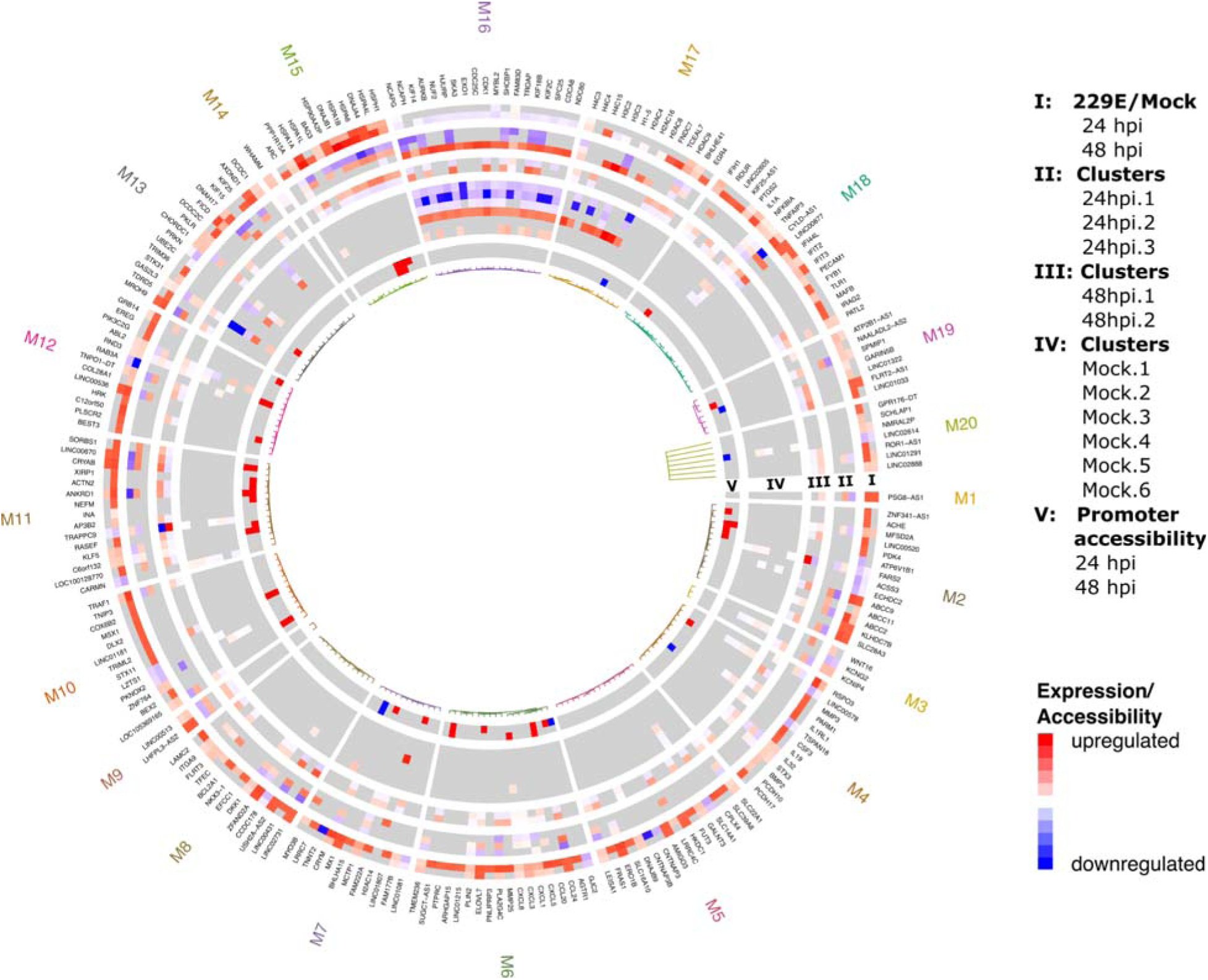
Mechanistic interpretation of transcriptomic responses to coronavirus 229E infection. MENTOR dendrogram and heatmap of genes of interest. From the outermost to the innermost rings, the heatmap tracks include differential expression levels in treatment versus control and 24 and 48 hpi (I), differential expression levels between cell clusters at 24 hpi (II), differential expression levels between cell clusters at 48 hpi (III), differential expression levels between control cell clusters (IV), and changes in promoter chromatin accessibility between treatment and control at 24 and 48 hpi (V).

## DISCUSSION

Single-cell analysis has advanced our understanding of immunity by revealing how individual cells in complex tissues respond to viral infection where heterogeneity often obscures specific cell contributions in bulk analyses. In contrast, cell culture models are often viewed as homogeneous systems of genetically identical cells presumed to respond uniformly to viral infection, an assumption that may overlook biologically meaningful cell-to-cell variability. Such variability may arise through multiple mechanisms, including differences in cell cycle state, metabolic activity, stochastic gene expression, or properties of the virus itself, such as the presence of defective viral genomes or viral haplotypes with different fitness^46,47^. Here, we provide insights from single-cell analyses of virus-infected cell cultures, uncovering fundamental mechanisms of antiviral defense and viral progression through distinctive transcriptional programs; these findings highlight that variability in infection outcomes likely reflects a combination of host-intrinsic and virus-driven factors, even within controlled *in vitro* systems.

In this study, we sought to determine the potential functional heterogeneity within ostensibly uniform cell cultures of human lung fibroblast cells (MRC-5) exposed to the common cold, human coronavirus (HCoV-229E). We hypothesized variability may arise either from differences in initial viral input and early infection dynamics, as well as subsequent differences in replication dynamics among neighboring cells, even under conditions of a defined multiplicity of infection, consistent with prior work showing that multiplicity of infection does not ensure uniform intracellular viral burden or response in influenza A^48^. Single-cell RNA-seq analysis at 24 and 48 hours post-infection (hpi) revealed substantial variation in viral load and host transcriptional responses, both within and between biological replicates even under controlled experimental conditions. These observations align with previous reports of similar variability in other host-pathogen viral systems^3^. Importantly, we also identified putative bystander cells with little to no detectable viral load, a phenomenon widely reported in single-cell infection studies and attributed to abortive infection or intrinsic resistance^7,16,49^. These cells either did not experience the infection, or successfully eliminated it, though we cannot definitively state which scenario is more likely. Indeed, our analysis is restricted to cells that survived infection (up to the sampling point), excluding direct assessment of the earliest responses in the most susceptible cells, any transient regulatory events, or the multiple iterations of viral replication and host feedback that have likely occurred by 24 hpi. Despite these caveats, the heterogeneity of viral read distributions indicates that infection and/or viral progression are not uniform across cells in culture.

In terms of cellular function, cells with higher viral read percentages upregulated heat shock protein genes and genes associated with unfolded protein responses, whereas cells with lower viral read percentages were enriched for cell cycle genes. While cell cycle state contributes to some heterogeneity, our analysis indicates that it does not fully explain the divergent transcriptomic responses observed in infected cells, which had reduced proliferative capacity and gene expression patterns associated with G_1_ arrest, consistent with prior reports that link coronaviruses to G_0/_G_1_ cell cycle arrest^50–52^. Cell-to-cell communication analysis further revealed that infection was associated with reduced ECM- and adhesion-related signaling, increased cytokine and growth factor communication, and reduced IL11-IL6ST interactions, consisten with decreased IL11-associated gp130 signaling. Further, this heterogeneity was closely linked with differential expression of NF-κB regulators and effectors, *NFKBIA*, *OTULIN*, *TNFAIP3*, and *PTGS2*, suggesting an important role for NF-κB regulation in shaping viral infection outcomes, in line with its well-established role as a central mediator of antiviral and inflammatory responses^53,54^. Supporting this, single-cell predictive expression networks (scPENs) revealed that NF-κB modulators are highly rewired upon infection, reinforcing the notion that viral gene expression intersects with the NF-κB signaling. At 24 hpi, highly rewired genes and genes with reduced connectivity were enriched for cell cycle processes, while genes gaining degree centrality were enriched for cell cycle checkpoint regulators, further supporting suppression of proliferation. From 24 to 48 hpi, genes losing global influence and network bridging roles were enriched for cell cycle processes, whereas genes gaining connectivity and influence over network information flow were enriched for stress and inflammatory responses, including the unfolded protein response, consistent with a systemic shift toward cellular stress and immune activation.

Finally, mechanistic network analysis using MENTOR revealed that gene modules associated with cell cycle processes, such as histone gene expression, mitotic control, and DNA repair, were suppressed in infected cells, while others linked NF-κB signaling to regulators of proliferation. Notably, modules enriched for transport, chemokine and cytokine signaling, cytoskeletal organization, and HSF1-mediated heat shock response contained genes whose promoter accessibility changed at 24 or 48 hpi, highlighting a key role for chromatin remodeling in orchestrating these transcriptional responses. Consistent with this, our bulk ATAC-seq analysis shows distinct changes in chromatin accessibility due to infection with markedly reduced global accessibility, consistent with previous findings of coronavirus-mediated impacts on chromatin remodeling^18,19^. Differentially accessible chromatin regions were enriched for binding motifs of NF-κB subunits p52 and p65/RelA at both 24 and 48 hpi, suggesting that NF-κB-driven transcriptional variability is shaped in part by chromatin accessibility dynamics. The essential role of NF-κB in coronavirus infections has been observed in other studies as well, where the severity of COVID-19 infection is associated with *NFKBIA* and *TNFAIP3*^55^, with the latter required for efficient HCoV-229E replication^18^. Additionally, HCoV-229E has been found to attenuate NF-κB activity via restriction of IKK (IκB kinase) complex activity. The persistence of promoter accessibility changes from 24 to 48 hpi further suggests that HCoV-229E infection induces stable, rather than transient, chromatin remodeling events that may shape later transcriptional states. Ultimately, while we cannot demonstrate the directionality or causality of the relationships between cell cycle, chromatin accessibility, or infection outcome, our results, alongside NF-κB’s known role in coordination of both immune responses and cell cycle progression, suggest that infection-induced variation in NF-κB activity shapes the transcriptional response in cultured lung fibroblasts. Future studies focused on targeted perturbation of NF-κB signaling or cell cycle regulators will further elucidate the mechanisms underlying the regulatory control of NF-κB on HCoV-229E viral infection progression.

By demonstrating how coronavirus infection remodels the host regulatory landscapes at the single-cell level, our findings advance our understanding of respiratory viral responses and offer insights relevant to antiviral research. More broadly, this work demonstrates that single-cell analysis of viral infection, even in a homogeneous cell culture, reveals mechanistic details that may be otherwise lost in bulk measurements. Future studies could extend these findings by examining regulatory dynamics at a finer temporal resolution in cell culture to capture transient events and responses in the most susceptible cells or deeper investigation into single-cell chromatin dynamics. Ultimately, clarifying how host regulatory states influence and shape infection outcome *in vitro* at the single-cell level may inform the development of more robust experimental models for antiviral research and help optimize the use of fibroblast lines in development of therapeutics.

## Availability of Data and Materials

All sequencing raw data generated during this study is available in the GEO repository with the following accession numbers: GSE310661 and GSE310663. (Tokens are available for reviewers.)

## Authors’ contributions

CRS and VV designed the experiments and profiling assays. VV, SA, and ES performed the viral exposures, cell harvest, and sample collection. VV performed the sample processing and preparation of single-cell RNA-seq and ATAC-seq libraries. All sequencing was performed at the LANL Genome Center. VV aligned and (re-)processed the scRNA-seq data and ATAC-seq data. CR processed ATAC-seq data. AV performed the scRNA-seq analysis, the downstream ATAC-seq analysis, Gene Ontology enrichment, MENTOR, and scPEN analyses and biological interpretation. KS developed MENTOR. AV and CRS wrote the manuscript and all authors edited the manuscript. CRS, DJ, & SS provided conceptualization, funding, and supervision for the project.

## Ethics Declarations

The authors declare no competing interests.

## SUPPLEMENTARY MATERIAL

**Figure S1.**
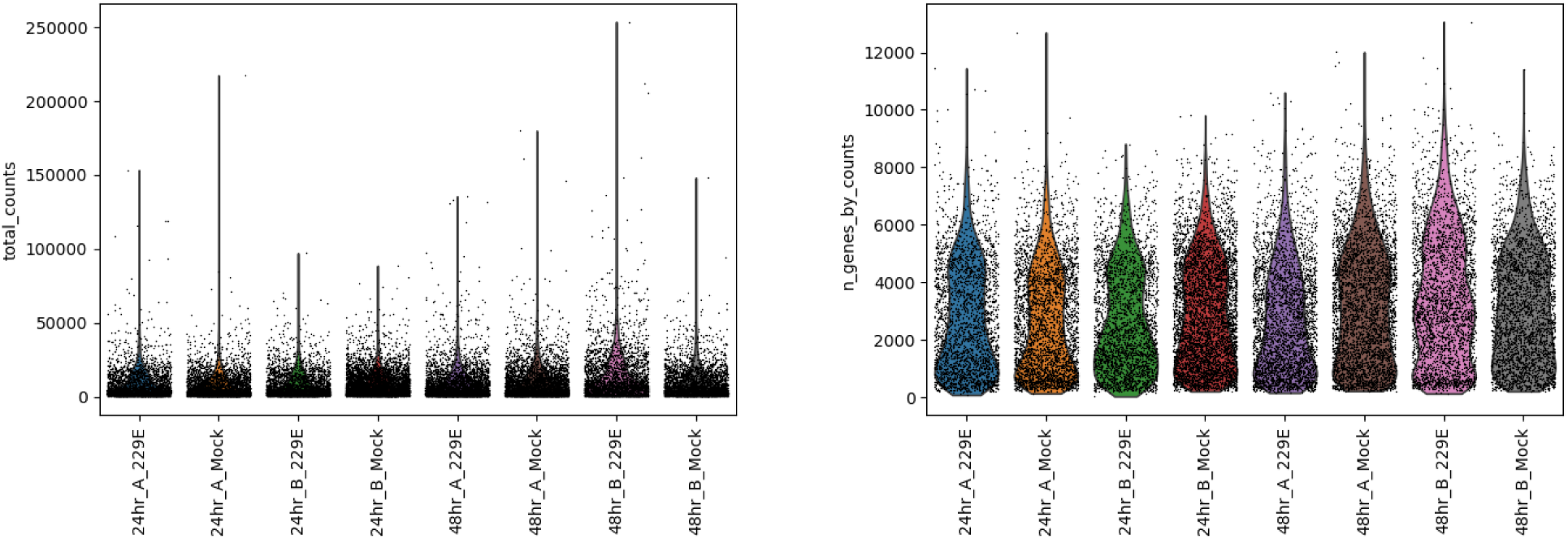
Quality control metrics per sample. **A)** the total counts per cell. B) the number of genes with at least 1 count per cell.

**Figure S2.**
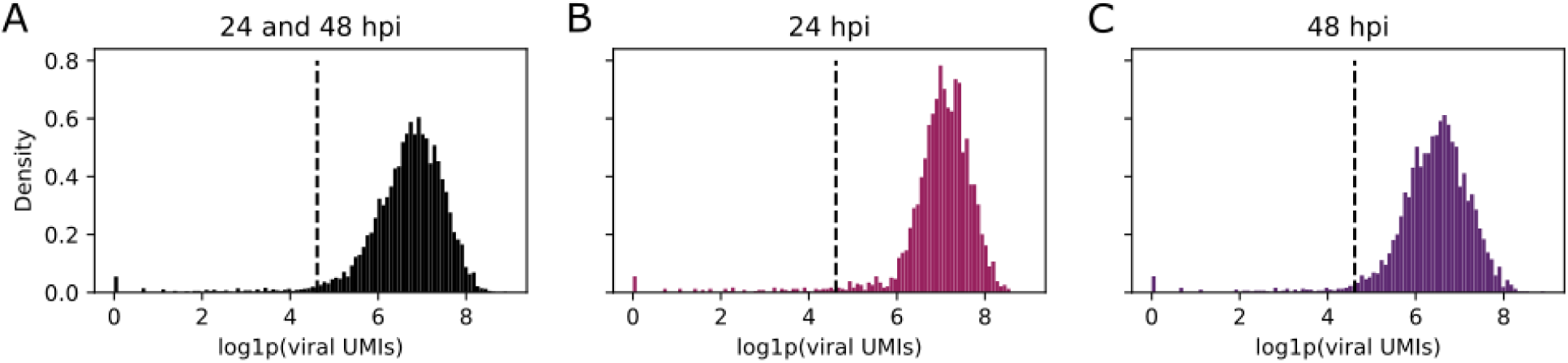
Viral read distributions in infected cells. **A)** Viral read distribution in all infected cells. **B)** Viral read distributions in infected cells at 24 hpi. **C)** Viral read distributions in infected cells at 48 hpi. Low-viral-load (putative bystander) cells are declared as cells with less than 100 UMIs as illustrated by the dashed line.

**Figure S3.**
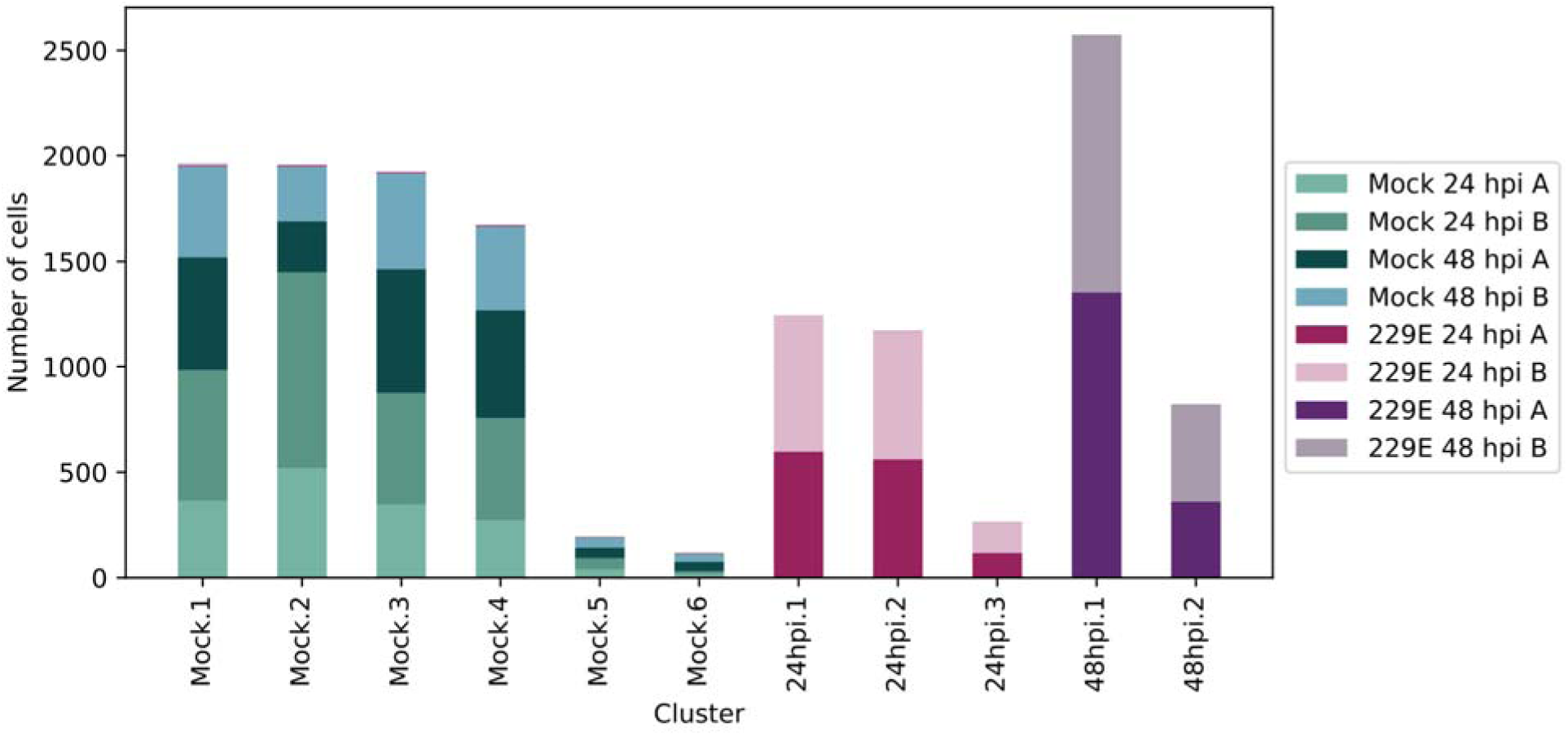
Sample contributions per cluster.

**Figure S4.**
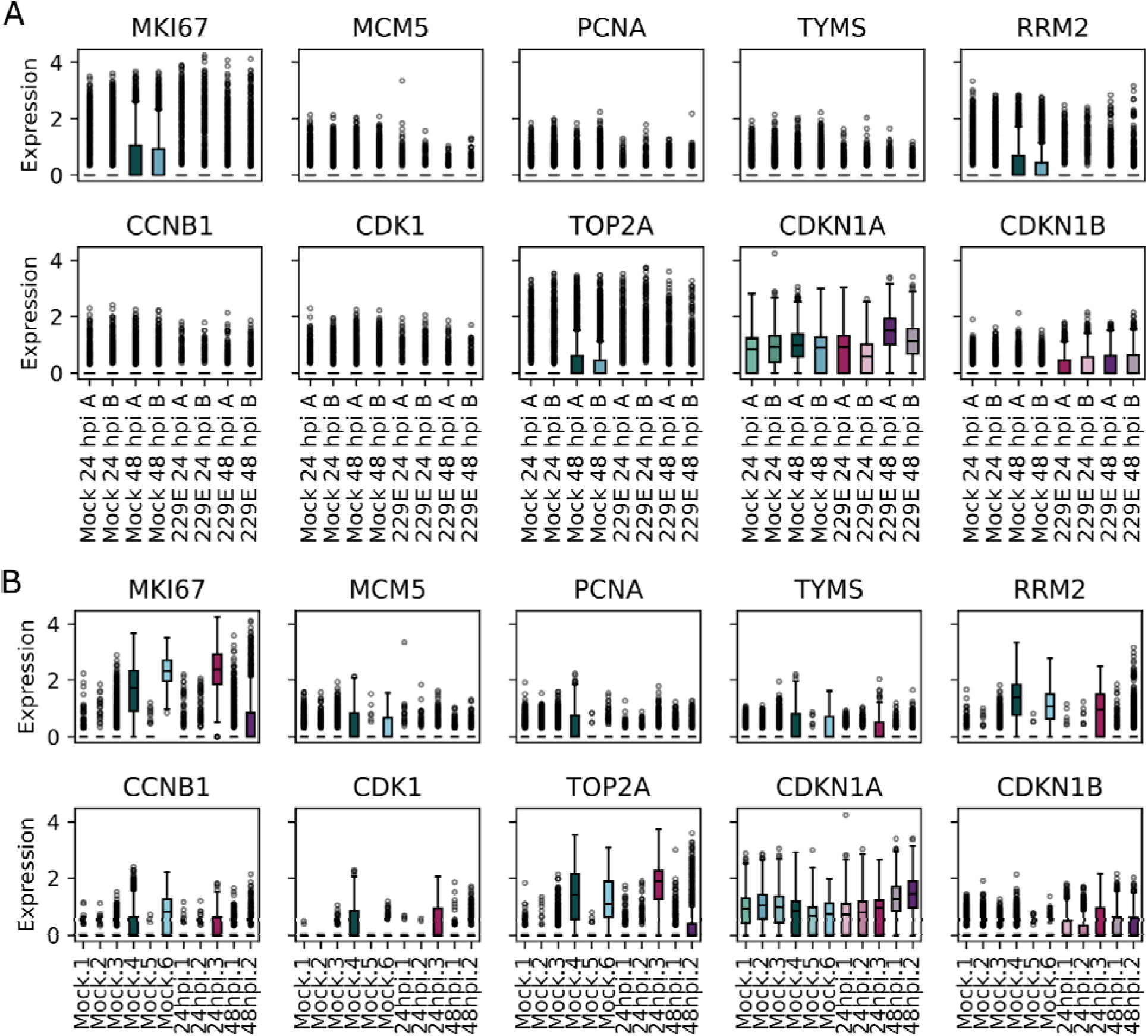
Cell cycle markers. **A)** Cell cycle markers expressed in each sample. **B)** Cell cycle markers expressed in each cluster. *MKI67* is a general proliferation marker; *MCM5*, *PCNA*, *TYMS*, and *RRM2* are S-phase markers; *CCNB1*, *CDK1*, and *TOP2A* are G_2_/M-phase markers; *CDKN1A* and *CDKN1B* are G_1_-arrest markers.

**Figure S5.**
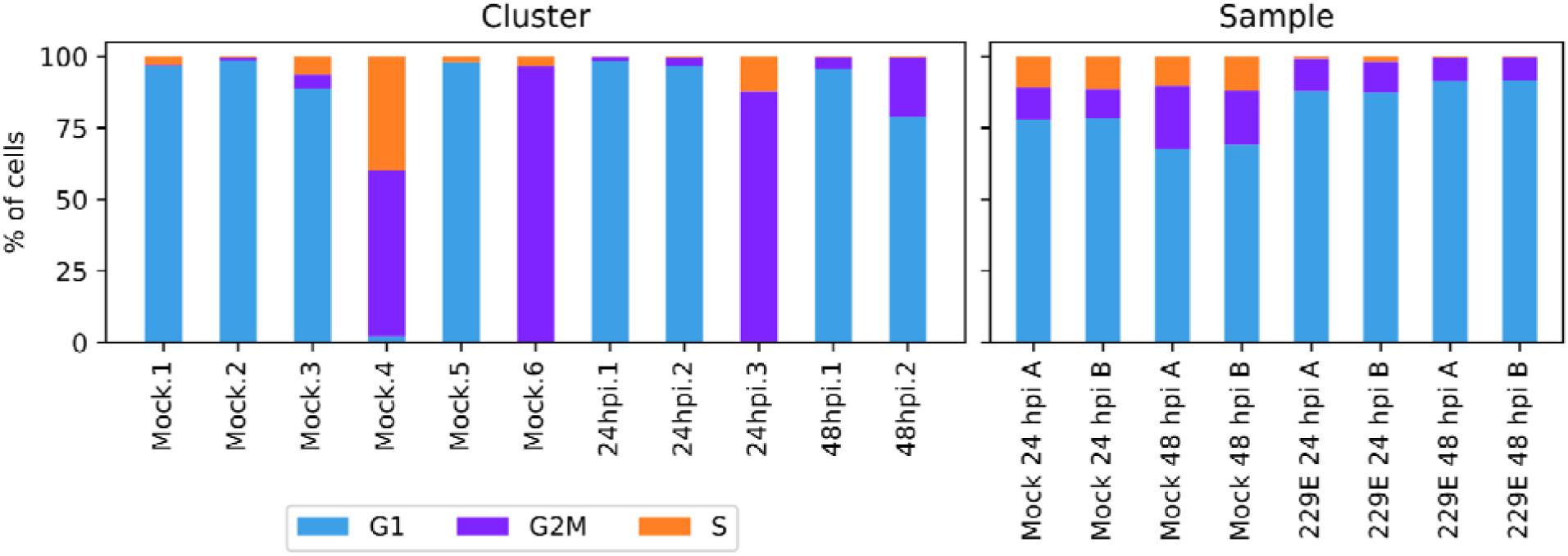
Percentage of cells in each cluster and sample that are in G_1_, G_2_/M, and S-phase.

